# Elucidating synergistic dependencies in lung adenocarcinoma by proteome-wide signaling-network analysis

**DOI:** 10.1101/289603

**Authors:** Mukesh Bansal, Jing He, Michael Peyton, Manjunath Kaustagi, Archana Iyer, Michael Comb, Michael White, John Minna, Andrea Califano

## Abstract

Signaling pathway models are largely based on the compilation of literature data from heterogeneous cellular contexts. Indeed, *de novo* reconstruction of signaling interactions from large-scale molecular profiling is still lagging, compared to similar efforts in transcriptional and protein-protein interaction networks. To address this challenge, we introduce a novel algorithm for the systematic inference of protein kinase pathways, and applied it to published mass spectrometry-based phosphotyrosine profile data from 250 lung adenocarcinoma (LUAD) samples. The resulting network includes 43 TKs and 415 inferred, LUAD-specific substrates, which were validated at >60% accuracy by SILAC assays, including “novel’ substrates of the EGFR and c-MET TKs, which play a critical oncogenic role in lung cancer. This systematic, data-driven model supported drug response prediction on an individual sample basis, including accurate prediction and validation of synergistic EGFR and c-MET inhibitor activity in cells lacking mutations in either gene, thus contributing to current precision oncology efforts.

## Introduction

Lung adenocarcinoma (LUAD) is a leading cause of cancer related deaths in United States, representing 40% of 225,500 new lung cancer cases every year, and has a 5-year survival rate of only 16 % (1). Excluding immunotherapeutic agents, which have recently shown significant success in a relatively small subset of patients (2), the most effective targeted therapies for this diseases were designed to inhibit tyrosine kinase proteins harboring genetic alterations that induce aberrant activation of downstream pathways (3-7). These the most frequent such actionable alterations include EGFR mutations and ALK-EML4 fusion events, in ~15% and ~3-7% of LUAD patients, respectively (8, 9). Yet, while targeted therapy is initially effective in a significant fraction of tumors harboring these genetic alterations, the vast majority of treated patients will either fail to respond or will develop resistance to mono-therapy (10, 11). In addition, most patient lack actionable alterations altogether. This suggests that novel approaches are critically needed.

A possible alternative to minimize emergence of resistance is combination therapy, a strategy that has been shown to be effective in many metastatic tumors, such as breast cancer and acute myeloid leukemia (12-14). However, systematic identification of effective drug combinations on a genetic alteration basis is difficult, because the number of patients presenting multiple actionable events is extremely low. As a result, combination therapy is generally hypothesized and tested on an empirical basis or based on elucidation of complex mechanisms of tumor cell adaptation. In addition, accurate prediction of response to available mono-therapy – including to EGFR inhibitors – in patients lacking any genetic alteration represents an equally relevant challenge, especially since a small fraction of EGFR^WT^ patients have been shown to respond to Afatinib, even though a predictive biomarker is not available. To address these limitations, we and other have proposed that rational design of combination therapy and the identification of critical targetable dependencies may require a more mechanistic and tumor-context-specific understanding of the molecular interactions that underlie their potential synergistic activity, starting with tyrosine kinases, which represent a critical class of pharmacological targets in cancer (15). Such an approach requires methodologies for the accurate and systematic elucidation of tumor-specific signaling transduction pathways.

Dissection of signal transduction networks represents a complex endeavor, requiring elucidation of hundreds of thousands of tissue-specific molecular interactions that mediate the post-translational modification of protein substrates. *In vitro* approaches generally fail to capture the tissue-specific nature of these interactions, thus providing “average” signal transduction pathways that are both incomplete and inaccurate. In addition, experimental approaches that have been successful in accelerating the analysis of molecular interactions in transcriptional regulation and protein-protein interaction in stable-complexes, such as those based on co-expression or yeast- 2-hybrid assays, do not easily translate to elucidating signaling interactions. Similarly, approaches based on the use of phospho-specific antibodies, while elegant and effective, are limited to only a handful of proteins. Computationally, compared to the many algorithms that have been developed for the reverse engineering of transcriptional and protein-complex interactions (16, 17), only a handful of experimentally validated algorithms are available for the dissection of signaling networks, none of which works at the proteome-wide level or is tumor-context specific (16, 18, 19).

Recent availability of proteome-wide molecular profile data, characterizing the abundance of phospho-tyrosine-enriched peptides by liquid chromatography coupled to tandem mass spectrometry (LC-MS/MS), suggests that additional methodologies may be developed to extend approaches that have been successfully applied to the dissection of transcriptional networks from gene expression profiling. In this manuscript, we propose extending the Algorithm for the Reconstruction of Accurate Cellular Networks (ARACNe) (20) for the reverse engineering of signal transduction networks from large-scale phosphoproteomic profiles. The new method, *pARACNe*, addresses critical issues that prevented the direct application of the original ARACNe algorithm on phosphoproteomic profile data. Briefly, the new algorithm addresses critical computational challenges presented by LC-MS/MS and spectral counting data, while incorporating enzymatic signaling characteristics into the algorithm design. In particular, pARACNE is designed to handle three critical issues resulting from the use of LC-MS/MS assays, including the highly sparse nature of phosphopeptide abundance data, the large amount of noise and missing data, and the degenerate peptides-to-protein mapping.

We applied pARACNE to the analysis of previously published, genome-wide phosphoproteomic data from 245 lung adenocarcinoma (LUAD) samples, including 151 fresh-frozen biopsies, 46 cell lines, as well as 48 normal lung tissues. The resulting network comprised 46 tyrosine kinases (TK) densely connected with 415 candidate substrates (including 377 proteins lacking any TK activity), representing the first genome-wide, tumor-context-specific model for a TK signal transduction network, capturing both protein-specific and phospho-site specific events. We validated substrate predictions for two “hubs,” whose activity may play a key role in determining sensitivity to Erlotinib and Crizotinib, two FDA-approved drugs for LUAD, including the EGFR and c-MET tyrosine kinases by independent SILAC assays and database analysis, with >60% accuracy. Of particular note, the inferred TK-substrate network provided unique information about tyrosine kinase auto-phosphorylation events, either direct (cis) or via a second kinase (trans).

Analysis of the resulting TK-network – by extending the VIPER (Virtual Proteomics by Enriched Regulon analysis) algorithm (21), an established method for the inference of Master Regulator proteins – recapitulated established genetic determinants of LUAD and was effective in predicting sensitivity to Erlotinib and Crizotinib combination therapy. Predicted sensitivities were validated in an independent set of LUAD cell lines, the majority of which harbored no genetic alterations in the corresponding genes. Furthermore, predictions based on the analysis of the corresponding patient cohort were strongly supported by genomic information, suggesting potential value in using these analyses for the identification of effective combination therapies in precision oncology.

## Results

### Overview of the pARACNe Algorithm

Enzymatic activity of tyrosine kinase (TK) proteins – as assessed by the ability to phosphorylate their downstream substrates – is effectively determined by their phosphorylated isoform abundance (**Fig. 1A, B**). Therefore, we reasoned that computational inference of TK-substrate interactions (**TK**→**S**) could be effectively performed by measuring dependencies between their respective phospho-states by mutual information analysis (22) over a large sample compendium **(Fig. 1C)**. Unfortunately, due to signal transduction cascade complexity and pathway cross-talk, such dependencies can manifest between protein pairs that are not involved in direct **TK**→**S** interactions. The ARACNe algorithm – previously designed for the reverse engineering of transcriptional networks – effectively addresses this problem by leveraging the Data Processing Inequality (23). This is a critical property of the mutual information that effectively allows disambiguating between direct and indirect interactions by assessing whether information transfer on any candidate direct interaction (e.g., **TK**_1_→**S**) is greater than transfer on every other indirect path (e.g., **TK**_1_→**TK**_2_→**S**). ARACNe has been highly successful in the experimentally validated dissection of transcriptional networks via analysis of large gene expression profile compendia. ARACNe-inferred targets of transcription factors were validated in multiple cellular contexts, with an accuracy of 70% to 80% (20, 24-27).

**Figure 1.**
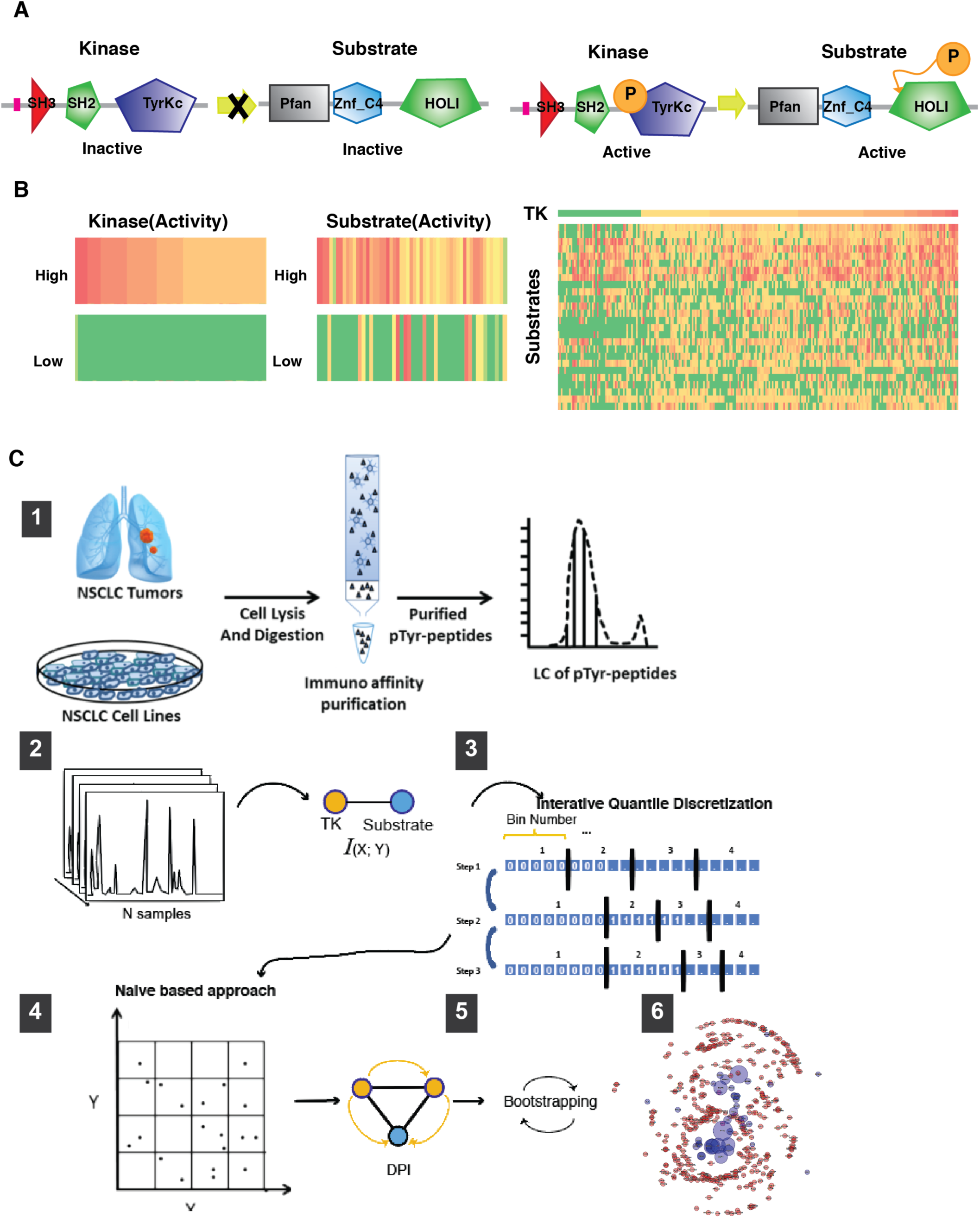
Framework for the reverse engineering of TK signaling networks from phosphoproteomic profiles. **(A)** Schematic diagram of a **TK**→**S** interaction. The non-phosphorylated kinase is inactive in terms of phosphorylating a substrate, while the active isoform successfully phosphorylates the substrate. **(B)** Schematic diagram showing the correlation between TK phosphorylation and that of its potential substrates. The first two rows in the heatmap show proteins representing candidate TK substrates **(C)** Illustration of the pARACNe framework including 6 steps. Step-1 depicts peptides collection from primary lung cancer tissue and cell lines for whole phosphrtyrosine proteomics quantification. Step-2 depicts inferences of TK→S interactions using Mutual Information by Step-3 Naïve-Bayes estimator and Step-4 of the iterative quantile discretization methods. Step-5 and 6 depict network pruning and bootstrapping to construct final network. **(C)**. Workflow of pARACNe from LC-MS/MS data normalization, IQD process, MI calculation, DPI process, bootstrapping to network consolidation.

However, ARACNe relies on molecular profile data that is both continuous and non-sparse, properties that are not always provided by quantitative proteomic data sets, which can be generated by a variety of methods. Those based on LC-MS/MS represent the most popular approaches (28), but different implementations have specific performance profiles in terms of analyte throughput, consistency of measurement of peptides across samples and linear dynamic range (29). Depending on the data acquisition method, one or both of these assumptions of ARACNe are violated in proteome-wide datasets generated by the most popular methods based on data-dependent acquisition. Particularly when employing quantification by spectral counting, as is typically conducted for global protein-protein interaction studies (30, 31), phosphoproteomic data is both discrete (i.e., generally represented by spectral counts) and very sparse, with a majority of peptides having zero spectral counts and presenting a significantly skewed distribution for low-abundance peptides.

To address these limitations, we developed a phospho-proteomic specific algorithm, pARACNe (phospho-ARACNe) (**Fig. 1C**), specifically designed to measure phospho-state dependencies between TKs and their candidate substrates from large-scale LC-MS/MS phosphoproteomic profiles. pARACNe thus extends the original ARACNe framework to allow systematic inference of **TK**→**S** interactions. Specifically, to handle the highly discrete nature of the data, we replaced the kernel-density and adaptive partitioning based mutual information estimators in the original algorithm with a bin-count based method (**Fig. 1C4**), using gold standard data to select the most effective number of bins [12] (see Methods). Furthermore, to deal with the skewed spectral count distribution, we introduce an iterative quantile discretization method, where samples are binned together, based on their spectral counts, to produce a distribution as close to uniform as possible (**Fig. 1C3,** Methods).

### pARACNe-inferred LUAD-specific TK-phosphorylation Network

We used pARACNe to reconstruct a LUAD-specific TK-signaling network, by analyzing phosphopeptide profiles obtained from 245 LUAD samples from Guo et al. (32). These data represent the abundance of peptides containing at least one phospho-tyrosine, as obtained by phosphoproteomic analysis of 46 LUAD cell lines, 151 LUAD tumors, and 48 adjacent normal samples. LC-MS/MS profiling produced spectral counts for 3,920 phospho-tyrosine containing peptides mapping to ~2,600 different proteins. Based on these data, pARACNE identified 2,611 candidate phospho-peptide/phospho-peptide dependencies, which could be further mapped to 2,064 unique **TK**→**S** interactions (**Table S1, S2)**. These represent interactions between 46 unique TKs and their candidate substrates. These include 174 **TK**_1_→**TK**_2_ interactions between two TKs (**Fig. 2A**), representing a statistically significant bias toward TK-TK interactions in the network (*p* = 10^−62^). This suggests that, within the complete TK signaling network, TKs themselves may form a more densely inter-connected subnetwork than previously assessed, providing potentially valuable novel information about adaptive response, pathway cross-talk, and auto-regulatory loops.

**Figure 2.**
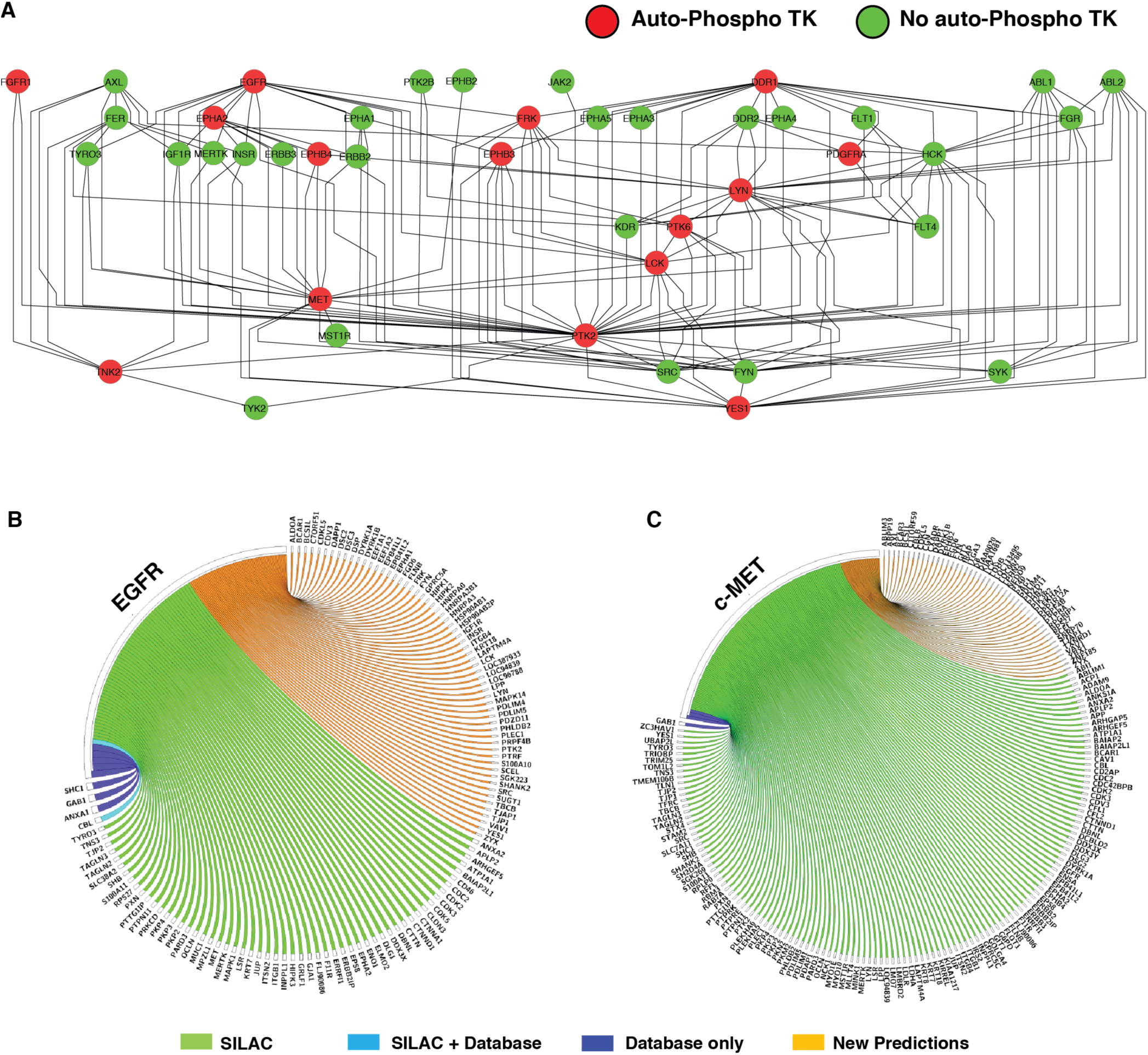
Predicted TK-TK network and validation of EGFR and c-MET prediction. **(A)** pARACNe-inferred densely inter-connected TK-TK network, with red nodes representing candidate TKs involved in auto-phosphorylation, where the phospho-state of a tyrosine is correlated with the phospho-state of a different tyrosine on the same TK protein. **(B)** pARACNe-inferred EGFR and **(C)** c-MET substrate overlap with SILAC-based and Database reported substrates, respectively.

Such highly interconnected structure provides potential functional advantage compared to less interconnected (i.e., “flat”) architectures, including the ability to provide more fine-grain response to a highly heterogeneous variety of exogenous signals and conditions, the ability to provide rapid adaptive response to changing stimuli, and the ability to preserve cell state via autoregulatory feedback. Consistent with the underlying biology, and in contrast to transcriptional networks, the vast majority of pARACNe-inferred interactions have a positive Spearman correlation, with higher counts of TK-mapped phosphopeptides corresponding to higher counts of candidate substrate-mapped ones. This is consistent with the fact that TKs only phosphorylate their substrates, thus inducing positive phospho-state correlation. Only a negligible number of inferred interactions (0.5%) were associated with a negative Spearman correlation (*N* = 11, *p ⩽* 0.05). These may represent either indirect interactions where the TK activates a substrate-specific phosphatase or direct interactions where phosphorylation of one phosphosite may prevent phosphorylation of another site on the same protein.

### LUAD Network Accuracy and Sensitivity Analysis

To estimate the accuracy of the inferred TK-signaling network, we investigated the substrates of two TK-proteins, EGFR and c-MET, representing high-affinity binding targets of existing FDA-approved TK inhibitors for LUAD. Specifically, we compared their pARACNe-inferred substrates to those reported in the phosphoDB database (33) and those supported by experimental evidence, based on previously published SILAC assays, following cell line treatment with associated, selective TK inhibitors. pARACNe inferred 123 EGFR substrates (**Fig. 2B**). Of these, 5 (blue and cyan) were included as high-confidence EGFR substrates in phosphoDB, out of 13 in total (38%), including the established EGFR auto-phosphorylation site. Moreover, 50 additional proteins (45%, green) showed significant decrease (at least 2 fold) in the abundance of their phosphorylated isoforms in SILAC assays (32), following treatment of H3255 cells with the EGFR inhibitor Gefitinib. Similarly, pARACNe predicted 179 c-MET substrates (**Fig. 2C)**. Notably, both of the established substrates reported in PhosphoDB were identified by pARACNE (100%, blue). Moreover, 126 additional proteins (71.5%, blue) showed significant decrease in the abundance of their phosphorylated isoforms in SILAC assays(32), following treatment of MKN45 cells with the first-generation c-MET-specific inhibitor Su11274.

We used MKN45 to assess overall prediction accuracy, even though it represents a gastric cancer cell line, because signaling networks should are much more conserved across tissue contexts than transcriptional ones. Indeed, while lineage-specific chromatin state represents a major determinant of transcriptional regulation, it only affects signal transduction in terms of overall protein availability. As a result, it is reasonable to expect that an even greater overlap of inferred vs. SILAC positive substrates may be achieved in native LUAD cells.

Taken together, these data suggest that pARACNe can identify a much larger subset of candidate substrates, while both identifying a significant proportion of established substrates (46% on average, based on phosphoDB) and maintaining high accuracy (~60% on average, by SILAC assays). This also suggest that, similar to transcription factor targets reported in the literature, TK substrates are still poorly characterized in existing repositories, even for highly relevant and exceedingly well-studied kinases such as EGFR and c-MET. As a result, pARACNe could provide significant novel hypotheses for TK→S interactions that can be validated as required. We should also note that the reported accuracy for pARACNe is estimated using SILAC data on a single cell line. SILAC assays have significant false negatives and it would be reasonable to expect that, once tested in additional cell lines, the accuracy of pARACNe could further increase. As a further performance benchmark, we used the same SILAC benchmark to test predictions by NetworkIN, a reverse engineering method based on protein sequence motif analysis and protein association networks (16). The analysis found almost no consensus with SILAC assays, with only one out of 33 NetworkIN-predicted EGFR substrate identified as significantly dephosphorylated following treatment with TK-specific inhibitors.

### Systematic, Network-based Inference of Pharmacological Dependencies

Once an accurate model of signal transduction in LUAD cells was established by pARACNe analysis, we interrogated the corresponding TK→S network using phosphoproteomic signatures from 46 LUAD cell lines to identify key dependencies for experimental validation. For this purpose, we extended the VIPER algorithm (Virtual Proteomics by Enriched Regulon analysis)(21), which was originally developed to identify the MR proteins that mechanistically regulate the transcriptional state of a tumor by assessing the enrichment of their transcriptional targets in differentially expressed genes in the tumor signature. VIPER and its predecessor MARINa (Master Regulator Inference algorithm) (34) have been instrumental in inferring MR proteins representing key functional determinants of tumor-related phenotypes in many cancer types, from glioblastoma (26, 27, 35), neuroblastoma (34), lymphoma (36, 37), and leukemia (38) to prostate (39-41) and breast adenocarcinoma(42-44), among others. We thus reasoned that VIPER could be modified to identify master regulator TKs, most likely to mechanistically regulate the differential phosphorylation pattern observed in a specific tumor sample (**Fig. 3A**). A specific additional value of the algorithm is that, as previously shown {Lefebvre, 2010 #1763}{Aytes, 2014 #2520}{Carro, 2010 #4021}, it could not only identify MR TK proteins, representing individual, pharmacologically accessible dependencies of the tumor, but also TKs representing potential synergistic MR-pair as candidate dependencies for combination therapy.

**Figure 3.**
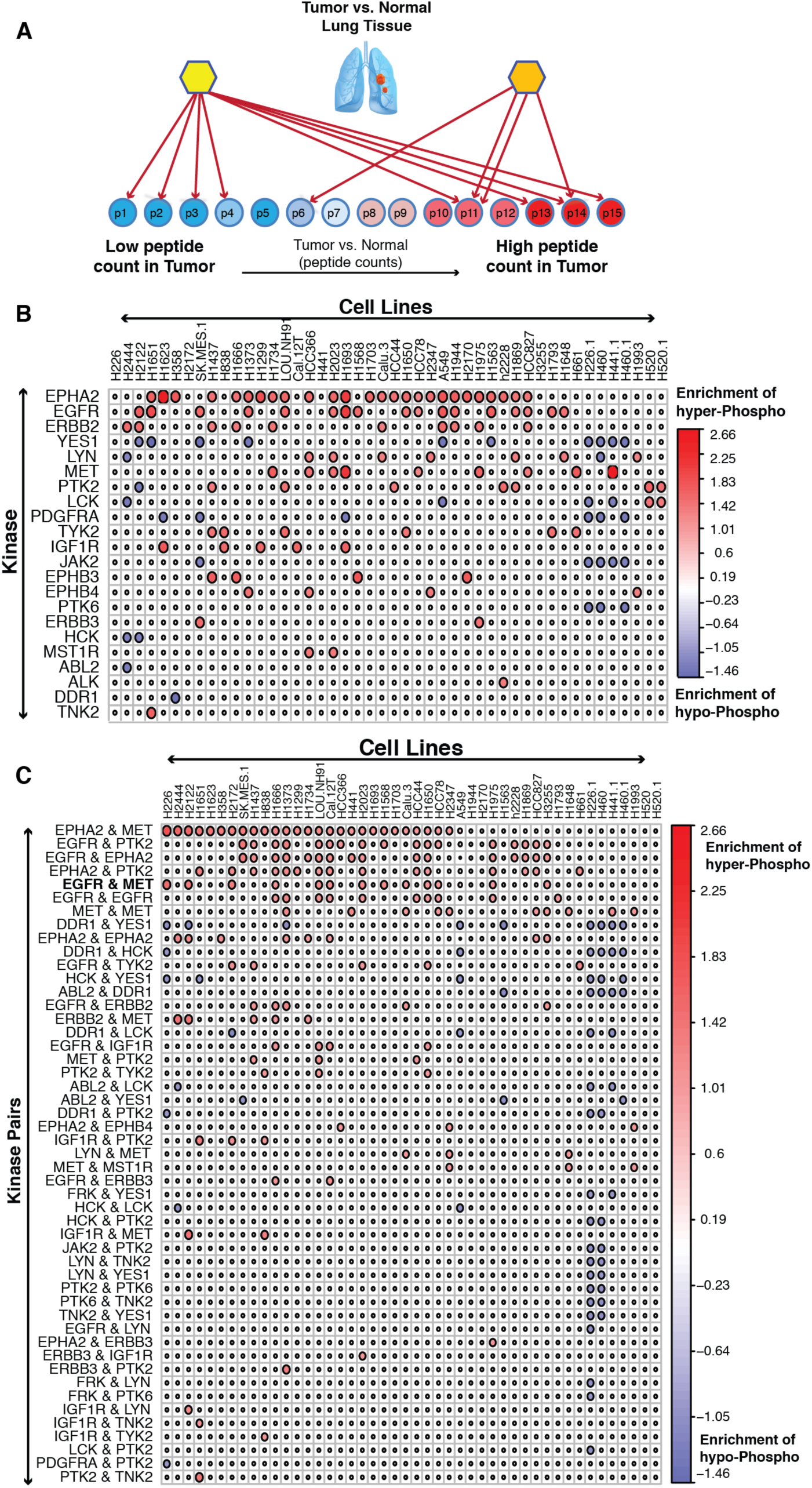
Inference of Master Regulator and combination. **(A)**. Schema of Master Regulator analysis in lung cancer using pVIPER. Prioritized Master Regulators **(B)** and Prioritized Master Regulator Pairs **(C)** as significantly activated (red circle) or de-activated (blue) in different lung cancer cell lines (column). Red color represents an enrichment of substrates hyper-phosphorylation by a Master Regulator or Master Regulator Pairs. Blue color represents that of hypo-phosphorylation.

VIPER can be easily modified to analyze phosphoproteomic signatures (pVIPER). Specifically, rather than assessing the enrichment of a protein’s transcriptional targets (regulon) in differentially expressed genes, pVIPER is designed to measure the enrichment of a TK’s substrates (signalon) in differentially phosphorylated proteins. Since inferred TK-substrate interactions are virtually all positive, this further increases the accuracy of the algorithm by supporting use of a single-tail enrichment analysis as opposed to the three-tail analysis of the original implementation. We first performed pVIPER analysis at the individual phosphopeptide level, rather than by averaging phosphopeptide state on a whole protein level. We then combined the result of the analysis across all phosphopeptides mapping to the same protein. Consistent with VIPER’s experimentally validated ability to identify synergistic master regulators proteins by transcriptional network analysis, pVIPER inferred several candidate synergistic TK interactions based on the statistical significance of the signature-enrichment of substrates shared by both TKs compared to that of substrates uniquely mapped to either one or the other TK (see Method section). Systematic VIPER analysis of phosphoproteomic profiles from 46 LUAD cell lines generated between 2 and 13 master regulator TKs or synergistic TK-pairs, as candidate pharmacologically actionable dependencies, for each cell line, thus generating a plausible number of hypothesis for each cell line (**Fig. 3B** and **Fig. 3C**).

### pVIPER Identifies LUAD-specific Dependencies

pVIPER analysis inferred several TK proteins as highly conserved individual dependencies across multiple cell lines, including the Ephrin type-A receptor 2 (*EPHA2*), epidermal growth factor receptor (*EGFR*), c-Met proto-oncogene (*MET*), and HER2 receptor tyrosine kinase 2 (*ERBB2*), suggesting a critical role of these proteins in the maintenance of LUAD cell line state. This is also in agreement with the functional role of these genes and the use of inhibitors of these kinases across a large panel of patients in multiple cancer types (45-49).

In contrast to these established LUAD cell line dependencies, we also identified several TKs as dependencies of specific cell lines. This can either be the result of associated genetic or epigenetic alterations in these cell lines or the result of field effects, where multiple genetic alterations or alterations in upstream pathway contribute to the cell line dependency on a specific TK activity. For instance, we identified ALK (Anaplastic Lymphoma Receptor Tyrosine Kinase) to be addiction point only in H2228 cell line. ALK is a conserved trans-membrane receptor tyrosine kinase (RTK) protein in the insulin-receptor super family. Chromosomal alterations involving *ALK* translocations and fusion events have been identified in several cancer types including LUAD (50, 51), diffuse large B-cell lymphomas (52), neuroblastoma (53), and inflammatory myofibroblastic tumors (54), among others. Additionally, ALK fusion events with other genes, including EML4 (Echinoderm Microtubule-associated protein Like 4) in LUAD lead to aberrant protein activity eliciting “oncogene addiction” (51). Presence of ALK-EML4 fusion transcripts, in ~3–7% of LUAD patients (55-57), is a strong predictor of response to ALK inhibitors, such as Crizotinib, among others (58, 59). Interestingly, among all available LUAD cell lines for which a phosphoproteomic profile was available, H2228 was the only one with an established ALK-EML4 fusion event and with established sensitivity to ALK inhibitor (60, 61). This further reflects the specificity of our analysis as this was the only cell line predicted to depend on ALK activity. Interestingly, we identified 4 additional H2228 dependencies, namely EGFR, Epha2, c-MET, and PTK2. H2228 sensitivity to EGFR inhibitors, in combination with ALK inhibitors, was already established (61).

### EGFR and c-MET are Predicted Dependencies in Multiple LUAD Cell Lines

As discussed, pVIPER analysis revealed several TK-pairs as candidate synergistic dependencies across several cell lines, such as Epha2/c-MET, EGFR/PTK2, EGFR/Epha2, Epha2/c-MET, and EGFR/c-MET. Among these the EGFR/c-MET pair emerged as the most conserved synergistic TK-pair across the available cell lines. In addition, EGFR and c-MET were also identified as candidate TK MRs in several of these cell lines, suggesting either a complementary or synergistic role for these proteins and a potential therapeutic opportunity for combination therapy in LUAD (62, 63).

To validate pVIPER-predicted, cell line specific EGFR/c-MET synthetic lethality, we selected a panel of 14 cell lines, 11 of which were predicted to be synergistically dependent on EGFR/c-MET (H226, H2122, H1666, H2172, Cal-12T, H2023, H1568, Calu-3, H1650, HCC78, and A549), as well as 3 negative controls with no predicted synergistic or individual dependencies on the two TKs (H2170, H460, and H520). To measure sensitivity to these agents, we used two different and complementary assays, including: (a) colony formation assay to assess long term sensitivity (**Fig. 4A and Methods**) and (b) 3-[4,5-dimethylthiazol-2-yl]-2,5 diphenyl tetrazolium bromide (MTS) assay for short term sensitivity analysis (**Fig. 5A, Table S3 and Methods**). For colony formation assays, cells were treated with either an EGFR inhibitor (Erlotinib, 1uM) or a c-MET/ALK inhibitor (Crizotinib, 0.1uM), either individually or in combination (see Methods). To evaluate synergistic dependency on EGFR/c-MET we used Excess Over Bliss (64), which measures the difference between the observed effect on colony formation and the effect expected from a purely additive model. For MTT assay, first, cells were treated with EGFR inhibitor (Erlotinib) or MET inhibitor (Crizotinib) individually at various concentrations to identify IC50 (concentration resulting in 50% cell death). Next, cells were treated with 1 uM of Erlotinib and varying concentrations of Crizotinib to identify combinations resulting in IC50 and used the combination index (CI) statistic (65) to measure interaction between the two drugs.

**Figure 4.**
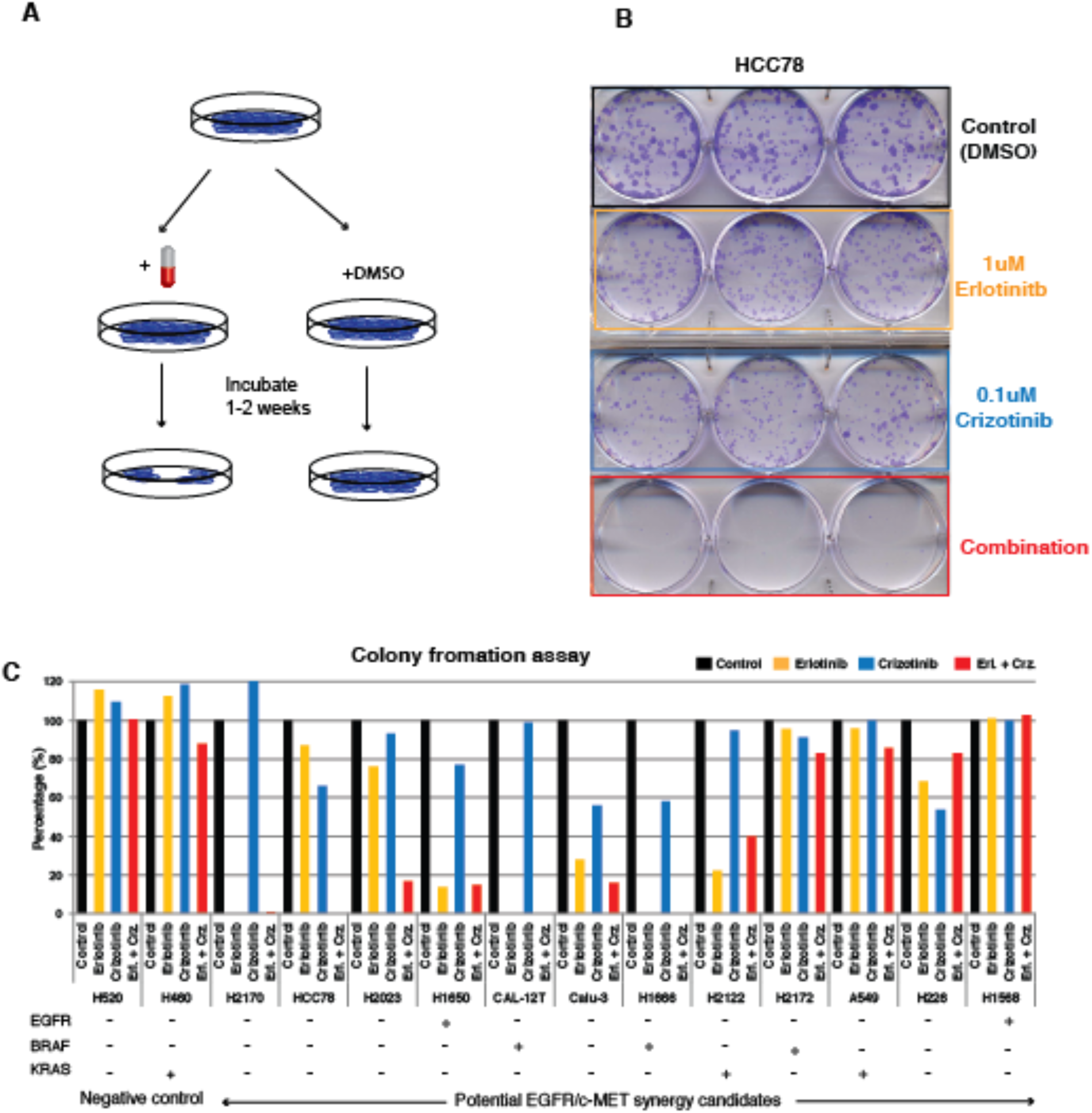
Experimental validation of EGFR and c-MET combination by colony formation assay. **(A)** Colony formation assay schema. **(B)** shows the image of long-term EGFR and c-MET double inhibition effects in HCC78 cell line with different treatments. **(C)** shows long-term clony formation data for 14 cell lines with different EGFR, BRAF and KRAS genomic mutation status.

**Figure 5.**
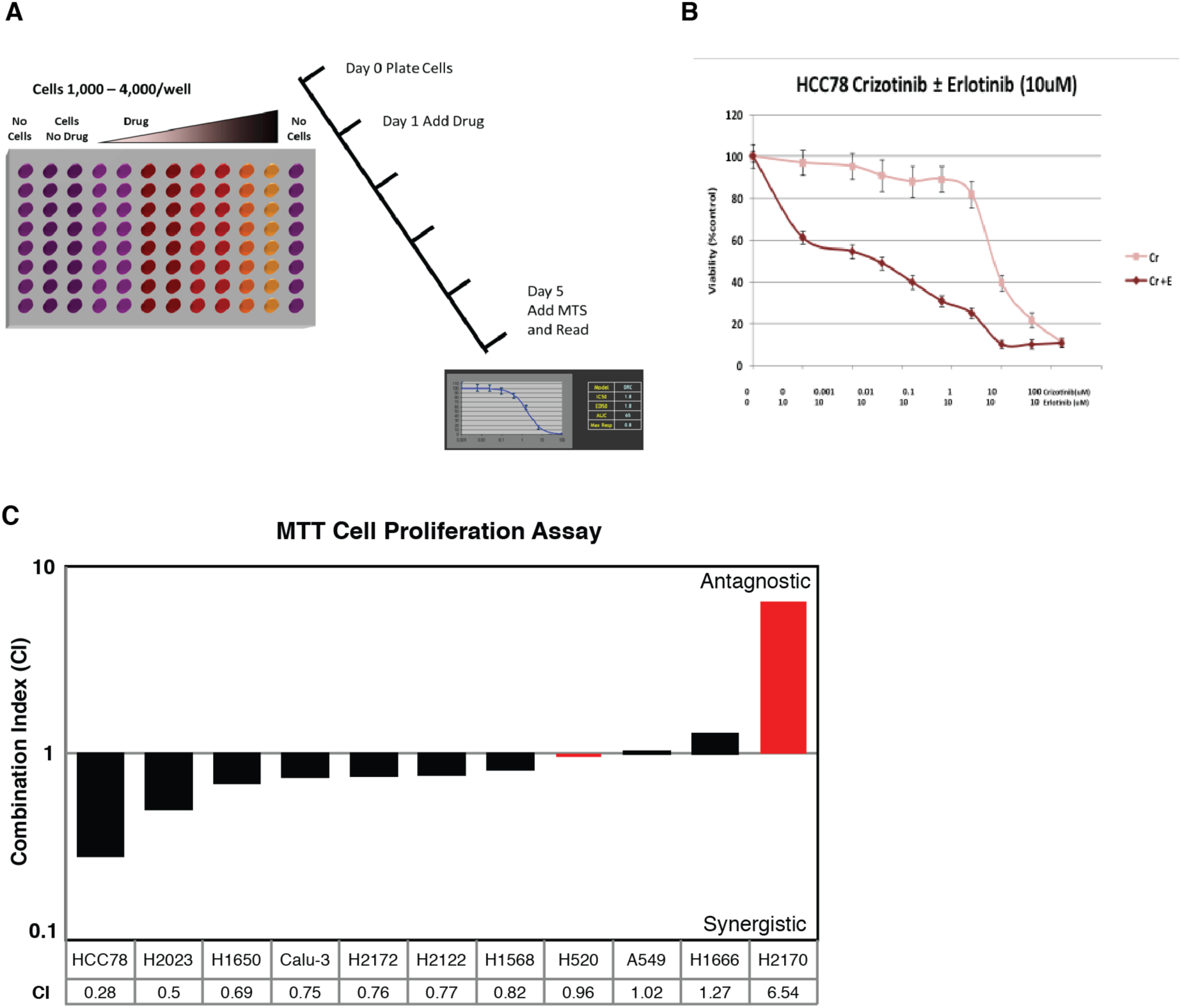
MTT Assay validation of EGFR and c-MET combination. **(A)**. MTT assay experimental schema. **(B)** MTT assay of HCC78 cell line shows synergistic effects of Crizotinib and Erlotinib treatment. **(C)** shows short-term effects of EGFR and c-MET inhibitors’ combination index in 11 cell lines include 2 control cell lines (red).

Across all 11 cell lines tested in colony formation assay, 8 showed significant sensitivity to either individual inhibitors (H226_E,C,_ H2122_E,_ H1666_E,C,_ Cal-12T_E,_ Calu-3_E,C,_ H1650_E_) or synergistic sensitivity to the combination (HCC78_E+C_ and H2023 _E+C_) (**Fig. 4B-C)**. Surprisingly, all of these cell lines were EGFR^WT^, ALK^WT^, and c-MET^WT^, except for H1650, which was EGFR^Mut^. Thus, based on standard of care criteria, 7 out of 8 cell lines would not have been considered as sensitive to either EGFR or ALK/c-MET inhibitors. Several cell lines presented striking sensitivity to either one (H2122_E_, Cal-12T_E,_ H1650_E_) or both inhibitors (H226_E,C,_ H1666_E,C,_ Calu-3_E,C_) in isolation, thus making the assessment of synergistic drug sensitivity difficult. In addition, three EGFR^WT^ cell lines harboring BRAF (Cal-12T and H1666) or KRAS (H2122) mutations were also highly sensitive to Erlotinib as a single agent, as predicted by pVIPER, despite the fact that KRAS pathway mutations are mutually exclusive with EGFR mutations and predictive of Erlotinib resistance (**Fig. 4C**). Finally, none of these cell lines was predicted to be sensitive to ALK inhibitors, suggesting that Crizotinib sensitivity is mediated by c-MET specific dependencies. Of the negative controls, only one (H2170) showed high sensitivity to Erlotinib. Taken together, 8/11 cell lines (73%) predicted as sensitive to the inhibitors were validated long term colony formation assays, while only 1/3 negative controls showed sensitivity to these agents (33%).

To evaluate the short-term interaction between EGFR and c-MET, we performed MTT assay across 11 cell lines (HCC78, H2023, H1650, Calu-3, H2172, H2122, H1568, A549, H1666, H520 and H2170) including 2 negative control cell lines (H520 and H2170). Similar to colony formation assay, we found synergistic sensitivity to EGFR and c-MET inhibitors in 6/9 cell lines (67%), with 5 cell lines showing strong synergy (CI ≤ 0.8) (**Fig. 5B-C**) and 1 borderline synergy (CI = 0.82), showing the consistency between two assays. However, for two cell lines, H1666 and H2170 (a negative control), results were inconsistent between long term colony formation and MTT assays. For both H1666 and H2170 cell lines, colony formation and MTT assay to showed sensitivity to Erlotinib alone, where colony formation assay has complete abrogation of colonies at 1 μM of Erlotinib, and later had *IC*_*50*_ =1.25 μM and 3.7 μM for H1666 and H2170 respectively. However, in combination therapy, MTT assay showed antagonism (CI >1), despite the fact that colony formation assay still showed complete abrogation which could be associated to the accumulation of new mutations in these cell lines. However, this is just hypotheses and needs to be verified by further experiments such as sequencing of these cell lines pre-and post-treatment.

### Phosphosite-specific Phosphorylation Predicts EGFR/c-MET Inhibitor Synergy

In previous section, we assessed the pVIPER predictions after consolidating the result at protein level. Following the results from MTT and colony formation assays, we reanalyzed the pVIPER predictions at the phosphopeptide level. Interestingly, this revealed that whenever synergistic EGFR/c-MET dependencies were predicted from phosphosite EGFR_1197_ and phosphosites other than c-MET1003 (H1666, Cal-12T, H1650), cell lines responded to Erlotinib in isolation, while when predictions were based on phosphosites EGFR_1197_ and c-MET_1003_ (HCC78, H2023, and Calu-3), cells exhibited bona fide synergistic sensitivity to the two inhibitors, with the only possible exception of Calu-3, which showed synergistic sensitivity in MTT assays and additive sensitivity to both inhibitors in colony formation assays. Conversely, when predictions were not based on either phosphotyrosine, cells exhibited no sensitivity to the individual inhibitors or the combination (H2172, H226, A549, H460, H520, H1568). Thus, predictions based on these two phosphosites produced no false positives (6 out of 6 predicted and validated as non-sensitive) and only 2 false negatives (H2170 and H2122), resulting in an error rate of only 2 out of 14 cell lines (14%, p = 0.0093 using fisher exact test).

This finding is in agreement with the established role of EGFR_1197_ as a predictor of EGFR inhibitor sensitivity (66). Intriguingly, when sensitivity was predicted using phosphosites other than EGFR_1197_ and c-MET_1003_, cell lines did not respond to the inhibitors, either individually or in combination. For these two peptides, we found their common substrates to be hyper phosphorylated in the sensitive cell lines **(Fig. 6A**) compared to the specific substrates of each of them, whereas cell line responding only to EGFR inhibitors showed more hyper phosphorylation of EGFR only substrates (**Fig. 6B**). Cell lines resistant to both EGFR and c-MET inhibitors either showed no change in the phosphorylation status or hypo-phosphorylation compared to the normal samples (**Fig. 6C**). Hence, either the common substrates of EGFR and c-MET, or the phosphorylation status of EGFR_1197_ and c-MET_1003_ could potentially be used as biomarkers for predicting therapy with the dual inhibitors. However, this conclusion is based on a very limited number of observations and lacks the statistical power. This finding needs a re-evaluation/validation using larger cohort of samples to establish an appropriate biomarker for combination therapy.

**Figure 6.**
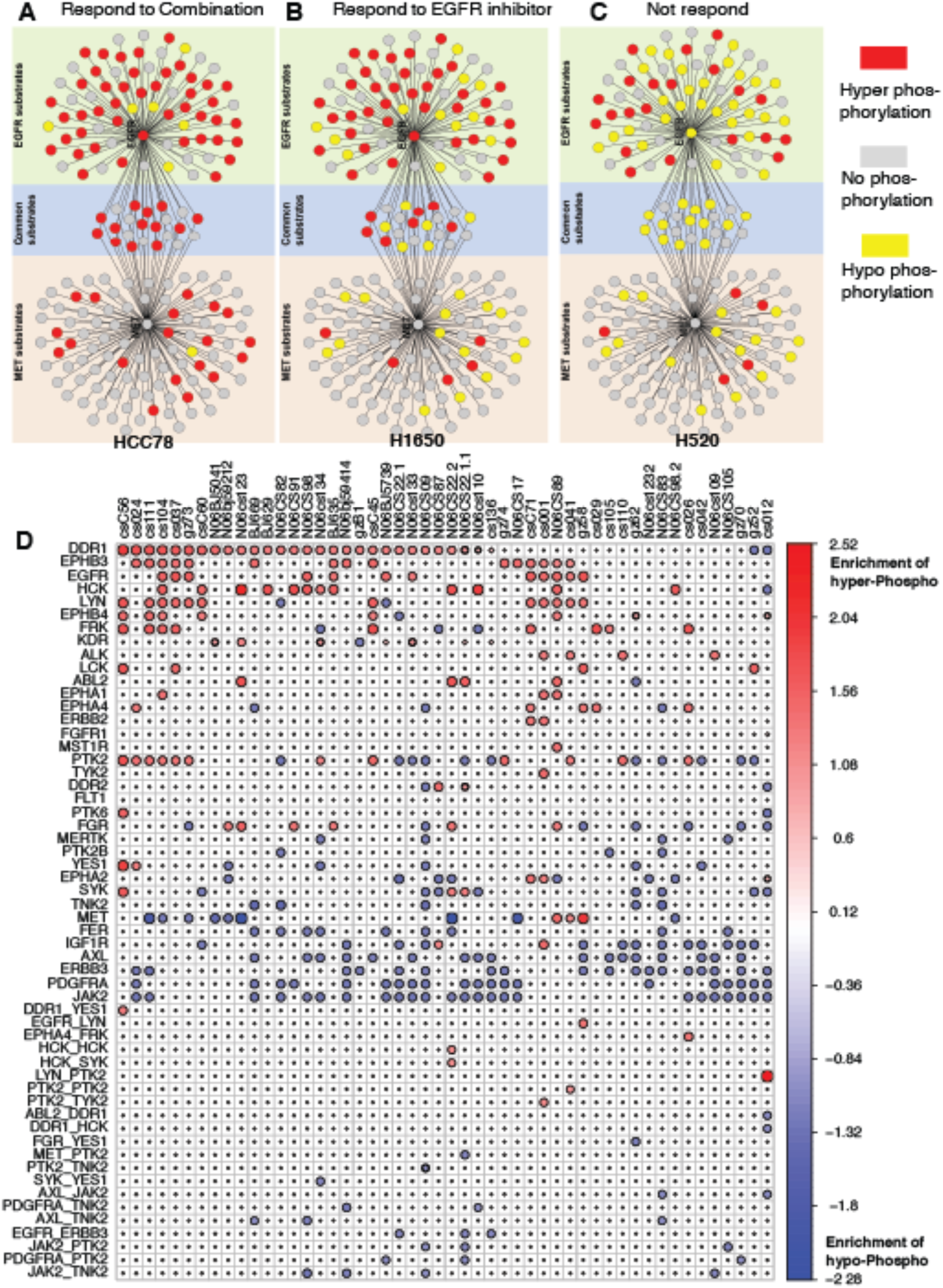
Master Regulating peptides in primary lung cancer samples EGFR and c-MET co-regulate in three scenarios. **(A)** when their common substrates are hyperphosphrylated, the patient responds to combination treatment well; **(B)** when most EGFR substrates are hyper-phosphorylated, the patient responds to EGFR inhibitor; **(C)** when substrates of both EGFR and c-MET are mostly hypophosphorylated, the patient does not respond. **(D)** show the Master Regulator and Master Regulator Pairs regulating hyper/hypo-phosphorylation of their network substrates in each primary samples.

### Systematic Inference of Patient-specific Dependencies

Similar to cell lines, when applied on patient data (32), pVIPER identified EGFR as one of the most common addiction points (**Fig. 6**). We inferred EGFR dependency in 12 patients. Of these, 5 harbored EGFR mutations, while the remaining 7 patients had not been tested for this mutation, showing a high consistency between our predictions and the genetic predisposition for sensitivity to EGFR inhibitors. In the entire cohort, there were only 3 patients with EGFR mutations that were not identified as EGFR dependent by pVIPER, resulting in an overall sensitivity of 62.5% (5/8). However, it is well known that >50% of patients harboring EGFR mutations do not respond to Erlotinib, suggesting that these may not be false negatives but rather patients with low activation of downstream EGFR pathways, despite their EGFR^Mut^ state. Similarly, our analysis identified candidate ALK dependencies in 4 patients. Of these one had an established TFG-ALK fusion, whereas the others had not been tested for ALK fusion events.

Across all patient samples, we observed Discoidin Domain Receptor-1 (DDR1) to be the most frequent addiction point, which was not predicted for any of the 46 cell lines. One reason for the difference is that DDR1 is collagen dependent and there may be differences in the 3D structure of the tumor and the cell lines growing on the plate. An independent study (67) in a cohort of 83 lung cancer specimens found that silencing of DDR1 in these samples leads to the hampering of cell survival, reduced invasiveness in collagen matrices, increased apoptosis in basal condition and decreased metastatic activity in model of tumor metastasis to bone, signifying it as a potential novel therapeutic target.

## Discussion

In this paper, we developed pARACNe to infer Tyrosine Kinase (TK) signaling network using published genome-wide phosphoproteomic data from lung cancer. The network prediction was validated using SILAC experiments, with high accuracy. Interrogation of the predicted TK-substrate network generated biologically meaningful hypotheses, followed by experimental validations illustrating the effectiveness of predicted kinase inhibitor combination, EGFR and c-MET combination inhibitors, in treating lung cancer cell lines. Furthermore, Master Regulator Analysis using patient proteomics data provides implications for using targeted agent combinations to treat patients based on their proteomic profile data.

Notably, pARACNe is significant and powerful as of its genome-wide scale and context-specificity in discovering global signaling cascading relationships, which were missing by previous methods. For example, methods proposed by Linding et al. (16) combine motif-based phospho-site predictions with information of physical association, co-occurrence, and co-expression to identify substrates with high specificity and accuracy, but with low coverage and lack of contextual specificity. Bender et al (17) used reverse phase protein assay data after various stimulations to cells and inferred signaling network using hidden Markov models and genetic algorithms. Even though the resulting networks are context specific, they lack genomic-scale coverage. There have been methods which used existing large-scale protein networks and prune them using transcriptomic information to identify signaling pathways (68-70). In addition, attempts have been made to reconstruct signaling network using gene expression data (71, 72). However, as signaling complexity lies mostly in upper level of cellular processes, inferring the cascades from downstream gene expression fails to capture all the dynamics. Also, PrePPI proposed by Zhang et al. (73) used protein structure-based methods to infer global protein-protein interaction, but this approach fails to address phosphorylation context specificity. Innovative uses of multiplex and microarray-based approaches, where multiple antibodies can be used to probe an ensemble of phosphoproteins, are finally becoming sufficiently mature to allow characterization of small pathways. Yet, these methods are still far from providing an unbiased, genome-wide view of signal-transduction processes and continue to be completely dependent on antibody specificity and availability. Similarly, assays developed specifically to monitor phosphorylation pathways, such as Stable Isotope Labeling with Amino acids in Cell culture (SILAC), provides a simple and straightforward approach to detect differential protein abundance. Coupled with phosphorylation enriched assays, it can provide high quality quantification for post-translation phosphorylation changes in cell lines. However, these methods are 1) laborious and costly; 2) can only be performed to dissect the substrates of a single enzyme at a time and 3) do not differentiate between direct and indirect targets.

To be noted, as the LC-MS/MS experiments used here was generated based on Tyrosine-kinase enrichment, which is only about ~2% of whole phosphoproteome. pARACNe is shown only on TK-substrates network. The current methodology could be extended to whole phosphoproteomic data based signaling network reconstruction where the data is available. In addition to label free based LC-MS/MS proteomics data used in this work, label based approaches, such as ITRAQ or TMT, could generate higher throughput whole proteomic profiles which might require future redesign of ARACNe to incorporate both kinases and phosphatases in regulating their downstream substrates. It is reasonable to expect that a version of ARACNe developed specifically to dissect signaling networks should work at least as well as its transcriptional counterpart. Since the relationship between the mRNA abundance of a gene encoding a transcription factor (TF) and the activity of the corresponding protein is much looser than that between the abundance of a phospho-isoform of a kinase and its enzymatic activity.

Even though research has attempted to identify addiction points based on gene expression data (74), predictions based on phosphoproteomic data appear superior in a way that they can directly reflect contextual specific signaling activity and are able to be directly targeted by kinase inhibitors. It is important to note that clinically, only patients with base-pair deletion at exon 19 (del746_A750) or a point mutation at exon 21 mutation (L858R) in EGFR shows sensitivity to EGFR inhibitor such as Cedirinib or Erlotinib (75).

## Acknowledgments

We thank George Rosenberger for manuscript advice. This work was supported by the National Cancer Institute (NCI) Cancer Target Discovery and Development program (1U01CA168426 to A.C., and 1U01CA176284 to JDM), NCI Research Centers for Cancer Systems Biology Consortium (1U54CA209997 to A.C.), NCI Outstanding Investigator Award (R35CA197745-02 to A.C.), NCI SPORE in Lung Cancer (P50CA70907 to JDM), and CPRIT (Award RP110708 to JDM).

## Author Contributions

M.B. and M.K. developed the algorithm. M.B. and J.H. performed the analysis. M.P. performed the experimental validation. A.I., M.B., M. W., J.M and A.C. guided the experiments. M.B. J.H., A.I. and A.C wrote the manuscript. M.B. and A.C. provided supervision and guidance. A.C. conceived, funded and administrated the project.

## Declaration of Interests

A.C.is a founder and shareholder of DarwinHealth Inc. and a member of the Tempus Inc. SAB and shareholder. Columbia University is a shareholder of DarwinHealth Inc.

## Methods and Data

### Phosphoproteomic Data

The previously published phosphoproteomic data used to reconstruct signaling network was download from (50). This dataset, representing the abundance of phospho-tyrosine containing peptides, was obtained by tandem mass spectrometry analysis of 46 non-small cell lung cancer (NSCLC) cell lines, 151 NSCLC tumors, and 48 normal lung tissue samples. Immunohistochemistry and a phospho-tyrosine specific antibody were used to screen 96 paraffin-embedded, formalin fixed tissue samples from NSCLC patients as described by Rikova et al.(50). About 30% of tumors showed high-levels of phospho-tyrosine expression. Immunoblotting of 46 NSCLC cell lines with a phospho-tyrosine specific antibody also showed heterogeneous reactivity especially in the molecular weight range characteristic of receptor tyrosine kinases.

Since phospho-tyrosine represents less than 1% of the cellular phosphoproteome, as determined by tandem mass spectrometry (MS/MS), and is difficult to analyze by conventional methods, immuno-affinity purification was performed with a phospho-tyrosine antibody to enrich for phospho-tyrosine containing peptides prior to tandem mass spectrometry. All tumors were identified as NSCLC based on standard pathology. Only those tumors with greater than 50% of cancer cells were considered for further analysis. NSCLC cell lines were grown overnight in low serum to reduce background phosphorylation from culture conditions.

Tandem MS profiling identified 3920 tyrosine phosphorylation sites on approximately 2600 different proteins. 85% of these sites appeared to be novel when compared against PhosphoSite (http://www.phosphosite.org), a comprehensive resource of known phosphorylation sites.

### pARACNe Algorithm

ARACNe is originally designed for gene expression data, where expression of genes is usually continuous and non-sparse. Quantitative data obtained from label-free LC-MS/MS by data-dependent acquisition via spectral counting is discrete and very sparse, with many phosphopeptides counts not observed for multiple peptides in each sample causing the current version of ARACNe to be not suitable for this data, which thus required major modifications to handle discrete data. To handle discrete abundances, we modified the mutual information computation approach from a kernel density estimation based method to a Naïve based estimation of mutual information, which is a histogram based technique(76). Briefly, consider a collection of *N* simultaneous measurements of two genes X and Y. Data is partitioned into M discrete bins *ai,* and *ki* denotes the number of measurements that lie within the bin *a*_*i*_. The probabilities *p*(*a*_*i*_) are then approximated by the corresponding relative frequencies of occurrence 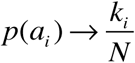 and the mutual information *I* (*X*,*Y*) between datasets X and Y is expressed as

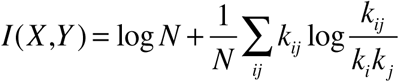

Here *k*_*ij*_ denotes the number of measurements where X lies in *a*_*i*_ and Y in *a*_*j*_ and *N* total number of samples.

Accuracy of mutual information is dependent on correct numbers of bins, M. To find the optimal number of bins we applied ARACNe on the whole dataset by varying M from 1 to 20 and testing the connections in predicted sub-network against the set of known connections (gold standard) from databases (phosphoDB) (33).

In case of continuous data, partitioning can be achieved by dividing the range of data into M equally spaced distance bins. Our data being discrete, equally spaced partitioning was not possible. So, to overcome this problem, we used an iterative approach of partitioning **(Fig. 1C, Fig. S1A)**. The basic idea is to divide the number of N data points into M with each bin containing equal number of data point. If the data point(s) with the same value falls into consecutive bin(s), we put those data point(s) into current bin and repartition the remaining points into remaining number of bins. We keep iterating this till we finish either the bins or there are no more data points to bin. For example, in **Fig. S1A**, we initially partition N points into 4 bins. The data points with 0 value does not fit into first bin and falls into subsequent bins, so we assign all data points with 0 value into first bin and repartition the remaining points into 3 bins. We keep on doing this process till we achieve 4 bins.

To evaluate initial performance and decide number of bins, we computed the network among all tyrosine kinases and substrates, parsed the sub-network between 49 tyrosine kinases and 114 substrates which were present in PhosphoSite database and compared the results with the connections present in database. From our analysis, we found that M=10 to be an optimal number **(Fig. S1B)** which gave us precision of 14% and sensitivity of 24%. This precision is an underestimate of real precision as in the gold standard many interactions are not present.

### Master Regulator Analysis

To discover the master regulator in various cell lines, we interrogated the network obtained from pARACNe using a novel algorithm, VIPER (Virtual Proteomics by Enriched Regulon analysis) (21), designed to infer kinases that are key players in a particular cell line. Protein activity is a good indicator of key kinases in a particular phenotype but often phosphorylated state of a protein is not sufficient to determine its activity both due to measurement noise in phosphorylated state as well as LC-MS/MS technique noise. To overcome this problem, VIPER infers kinase activity from the global kinase substrate relationship and its biological relevance by overlapping this information in a particular phenotype-specific program.

VIPER requires a network model and signature of the phenotype transition (i.e., all genes ranked by their differential phosphorylation in two phenotypes). Here, the signature, *S*_*kin*_, was obtained by t-test analysis by comparing each cell line against all normal samples. First, we associate each kinase with positive and negative activity targets, by computing the correlation between each kinase and its predicted substrates and selecting only those substrates which had a significant correlation (p-value ≤ 0.05, Bonferroni corrected). Second, for each kinase we computed an *activity* by measuring the enrichment of the *S*_*kin*_ signature in predicted substrates list, separately for both positive and negative correlated, (*S_kin_-enrichment*). Enrichment was computed by Gene Set Enrichment Analysis (GSEA). Since very small percentage of kinases are found to have negative correlation, we did not use those interactions to evaluate enrichment.

### Cell Culture

All cell lines were grown in RPMI-1640 with 5% fetal bovine serum and incubated at 37°C in a humidified atmosphere containing 5% CO2. Cell lines were fingerprinted using the Perplex 1.2 system (Promega, Madison, WI). Fingerprints were compared to those generated at ATCC and/or our internal database.

### MTS Assays

Short term MTS assays (CellTiter 96^®^ AQ_ueous_ One Solution Cell Proliferation Assay, Promega, Madison WI) were performed as previously described in (77). Specifically, each drug concentration is octuplicated and the mean with standard deviation of all replicates were used to generate a curve to allow calculation of the drug IC50 (Inhibitory Concentration of 50%) value. The assays were repeated at least 3 times and the IC50s are the average of all replicates.

### Colony-formation Assays

Long term colony formation assays were performed in triplicate in 6-well plates. Cells were added to media containing drug and incubated for 1-2 weeks such that control wells (no drug) contained colonies of 50-70 cells each. At such time media was removed and all wells stained with a solution containing 0.5% crystal violet and 6% glutaraldehyde for 1 hour. The plates were then rinsed, dried, and colonies were manually counted.

### SILAC Experiments

EGFR SILAC experiment was performed in H3255 cell line by treating samples with Gefitinib. c-MET SILAC experiment was performed in c-MET-driven gastric cancer cell line, MKN45, by using c-MET inhibitor Su11274. For both genes, cells were treated with inhibitors for 3 and 24hr. For control, cells were grown in same conditions but were not treated with the drug. For our comparison we combined the peptides, which were differentially obtained between treated and untreated samples, for 3 and 24 hr. More details about the experiment can be obtained from Guo et al (32).

#### Gold standard

In PhosphoSite database, there were 282 connections between 49 tyrosine kinases and 114 substrates.

## Supplemental Information

***Fig. S1. Performance of the pARACNe Algorithm***.

**(A).** To select optimal bin number in pARACNe algorithm, precision and recall curves for various number of bins were computed. Black curve is when no binning of data is done. When using 10 bins, the algorithm achieved the best performance.

***Table S1. pARACNe-inferred TK-peptides/substrate-peptides Interaction Network***.

***Table S2. pARACNe-inferred TK-Protein/Substrate Interaction Network***.

***Table S3. Colony Formation Assay and MTS Assay Results***.

## References

1. Ettinger DS, Akerley W, Borghaei H, Chang AC, Cheney RT, Chirieac LR, et al. Non-small cell lung cancer. Journal of the National Comprehensive Cancer Network : JNCCN. 2012;10(10):1236–71.

2. Karasaki T, Nagayama K, Kawashima M, Hiyama N, Murayama T, Kuwano H, et al. Identification of Individual Cancer-Specific Somatic Mutations for Neoantigen-Based Immunotherapy of Lung Cancer. J Thorac Oncol. 2016;11(3):324–33.

3. Pasqualucci L, Kitaura Y, Gu H, Dalla-Favera R. PKA-mediated phosphorylation regulates the function of activation-induced deaminase (AID) in B cells. Proc Natl Acad Sci U S A. 2006;103(2):395–400.

4. Fiedler D, Braberg H, Mehta M, Chechik G, Cagney G, Mukherjee P, et al. Functional organization of the S. cerevisiae phosphorylation network. Cell. 2009;136(5):952–63.

5. Gainor JF, Varghese AM, Ou SH, Kabraji S, Awad MM, Katayama R, et al. ALK rearrangements are mutually exclusive with mutations in EGFR or KRAS: an analysis of 1,683 patients with non-small cell lung cancer. Clin Cancer Res. 2013;19(15):4273–81.

6. Janne PA, Shaw AT, Pereira JR, Jeannin G, Vansteenkiste J, Barrios C, et al. Selumetinib plus docetaxel for KRAS-mutant advanced non-small-cell lung cancer: a randomised, multicentre, placebo-controlled, phase 2 study. Lancet Oncol. 2013;14(1):38–47.

7. Mak KS, Gainor JF, Niemierko A, Oh KS, Willers H, Choi NC, et al. Significance of targeted therapy and genetic alterations in EGFR, ALK, or KRAS on survival in patients with non-small cell lung cancer treated with radiotherapy for brain metastases. Neuro Oncol. 2014.

8. Vendrell JA, Taviaux S, Beganton B, Godreuil S, Audran P, Grand D, et al. Detection of known and novel ALK fusion transcripts in lung cancer patients using next-generation sequencing approaches. Sci Rep. 2017;7(1):12510.

9. Peters S, Camidge DR, Shaw AT, Gadgeel S, Ahn JS, Kim DW, et al. Alectinib versus Crizotinib in Untreated ALK-Positive Non-Small-Cell Lung Cancer. N Engl J Med. 2017;377(9):829–38.

10. Tsvetkova E, Goss GD. Drug resistance and its significance for treatment decisions in non-small-cell lung cancer. Curr Oncol. 2012;19(Suppl 1):S45–51.

11. Wilson FH, Johannessen CM, Piccioni F, Tamayo P, Kim JW, Van Allen EM, et al. A functional landscape of resistance to ALK inhibition in lung cancer. Cancer Cell. 2015;27(3):397–408.

12. O'Shaughnessy J, Miles D, Vukelja S, Moiseyenko V, Ayoub JP, Cervantes G, et al. Superior survival with capecitabine plus docetaxel combination therapy in anthracycline-pretreated patients with advanced breast cancer: phase III trial results. J Clin Oncol. 2002;20(12):2812–23.

13. Pegram MD, Slamon DJ. Combination therapy with trastuzumab (Herceptin) and cisplatin for chemoresistant metastatic breast cancer: evidence for receptor-enhanced chemosensitivity. Semin Oncol. 1999;26(4 Suppl 12):89–95.

14. Ravandi F, Cortes JE, Jones D, Faderl S, Garcia-Manero G, Konopleva MY, et al. Phase I/II study of combination therapy with sorafenib, idarubicin, and cytarabine in younger patients with acute myeloid leukemia. J Clin Oncol. 2010;28(11):1856–62.

15. Wu P, Nielsen TE, Clausen MH. Small-molecule kinase inhibitors: an analysis of FDA-approved drugs. Drug Discov Today. 2016;21(1):5–10.

16. Linding R, Jensen LJ, Ostheimer GJ, van Vugt MA, Jorgensen C, Miron IM, et al. Systematic discovery of in vivo phosphorylation networks. Cell. 2007;129(7):1415–26.

17. Bender C, Henjes F, Frohlich H, Wiemann S, Korf U, Beissbarth T. Dynamic deterministic effects propagation networks: learning signalling pathways from longitudinal protein array data. Bioinformatics. 2010;26(18):i596–602.

18. Papin JA, Hunter T, Palsson BO, Subramaniam S. Reconstruction of cellular signalling networks and analysis of their properties. Nature reviews. 2005;6(2):99–111.

19. Hashemikhabir S, Ayaz ES, Kavurucu Y, Can T, Kahveci T. Large-scale signaling network reconstruction. IEEE/ACM Trans Comput Biol Bioinform. 2012;9(6):1696–708.

20. Basso K, Margolin AA, Stolovitzky G, Klein U, Dalla-Favera R, Califano A. Reverse engineering of regulatory networks in human B cells. Nat Genet. 2005;37(4):382–90.

21. Alvarez MJ, Shen Y, Giorgi FM, Lachmann A, Ding BB, Ye BH, et al. Functional characterization of somatic mutations in cancer using network-based inference of protein activity. Nat Genet. 2016;48(8):838-47.

22. MacKay DJC. Information theory, inference, and learning algorithms. Cambridge, UK ; New York: Cambridge University Press; 2003. xii, 628 p. p.

23. Margolin AA, Nemenman I, Basso K, Wiggins C, Stolovitzky G, Dalla Favera R, et al. ARACNE: an algorithm for the reconstruction of gene regulatory networks in a mammalian cellular context. BMC bioinformatics. 2006;7 Suppl 1:S7.

24. Basso K, Saito M, Sumazin P, Margolin AA, Wang K, Lim WK, et al. Integrated biochemical and computational approach identifies BCL6 direct target genes controlling multiple pathways in normal germinal center B cells. Blood. 2010;115(5):975–84.

25. Della Gatta G, Palomero T, Perez-Garcia A, Ambesi-Impiombato A, Bansal M, Carpenter ZW, et al. Reverse engineering of TLX oncogenic transcriptional networks identifies RUNX1 as tumor suppressor in T-ALL. Nat Med. 2012;18(3):436–40.

26. Carro MS, Lim WK, Alvarez MJ, Bollo RJ, Zhao X, Snyder EY, et al. The transcriptional network for mesenchymal transformation of brain tumours. Nature. 2010;463(7279):318–25.

27. Sonabend AM, Bansal M, Guarnieri P, Lei L, Amendolara B, Soderquist C, et al. The transcriptional regulatory network of proneural glioma determines the genetic alterations selected during tumor progression. Cancer Res. 2014;74(5):1440–51.

28. Aebersold R, Mann M. Mass-spectrometric exploration of proteome structure and function. Nature. 2016;537(7620):347–55.

29. Gillet LC, Leitner A, Aebersold R. Mass Spectrometry Applied to Bottom-Up Proteomics: Entering the High-Throughput Era for Hypothesis Testing. Annu Rev Anal Chem (Palo Alto Calif). 2016;9(1):449–72.

30. Huttlin EL, Ting L, Bruckner RJ, Gebreab F, Gygi MP, Szpyt J, et al. The BioPlex Network: A Systematic Exploration of the Human Interactome. Cell. 2015;162(2):425–40.

31. Huttlin EL, Bruckner RJ, Paulo JA, Cannon JR, Ting L, Baltier K, et al. Architecture of the human interactome defines protein communities and disease networks. Nature. 2017;545(7655):505–9.

32. Guo A, Villen J, Kornhauser J, Lee KA, Stokes MP, Rikova K, et al. Signaling networks assembled by oncogenic EGFR and c-Met. Proc Natl Acad Sci U S A. 2008;105(2):692–7.

33. Giansanti P, Aye TT, van den Toorn H, Peng M, van Breukelen B, Heck AJ. An Augmented Multiple-Protease-Based Human Phosphopeptide Atlas. Cell Rep. 2015;11(11):1834–43.

34. Lefebvre C, Rajbhandari P, Alvarez MJ, Bandaru P, Lim WK, Sato M, et al. A human B-cell interactome identifies MYB and FOXM1 as master regulators of proliferation in germinal centers. Mol Syst Biol. 2010;6:377.

35. Chudnovsky Y, Kim D, Zheng SY, Whyte WA, Bansal M, Bray MA, et al. ZFHX4 Interacts with the NuRD Core Member CHD4 and Regulates the Glioblastoma Tumor-Initiating Cell State. Cell Rep. 2014;6(2):313–24.

36. Ying CY, Dominguez-Sola D, Fabi M, Lorenz IC, Hussein S, Bansal M, et al. MEF2B mutations lead to deregulated expression of the oncogene BCL6 in diffuse large B cell lymphoma. Nat Immunol. 2013;14(10):1084-+.

37. Bisikirska B, Bansal M, Shen Y, Teruya-Feldstein J, Chaganti R, Califano A. Elucidation and Pharmacological Targeting of Novel Molecular Drivers of Follicular Lymphoma Progression. Cancer Res. 2016;76(3):664–74.

38. Piovan E, Yu J, Tosello V, Herranz D, Ambesi-Impiombato A, Da Silva AC, et al. Direct reversal of glucocorticoid resistance by AKT inhibition in acute lymphoblastic leukemia. Cancer Cell. 2013;24(6):766–76.

39. Aytes A, Mitrofanova A, Lefebvre C, Alvarez MJ, Castillo-Martin M, Zheng T, et al. Cross-species regulatory network analysis identifies a synergistic interaction between FOXM1 and CENPF that drives prostate cancer malignancy. Cancer Cell. 2014;25(5):638–51.

40. Mitrofanova A, Aytes A, Shen C, Abate-Shen C, Califano A. A systems biology approach to predict drug response for human prostate cancer based on in vivo preclinical analyses of mouse models. Cell Reports. 2015;12:1–12.

41. Talos F, Mitrofanova A, Bergren SK, Califano A, Shen MM. A computational systems approach identifies synergistic specification genes that facilitate lineage conversion to prostate tissue. Nature communications. 2017;8:14662.

42. Walsh LA, Alvarez MJ, Sabio EY, Reyngold M, Makarov V, Mukherjee S, et al. An Integrated Systems Biology Approach Identifies TRIM25 as a Key Determinant of Breast Cancer Metastasis. Cell Rep. 2017;20(7):1623–40.

43. Rodriguez-Barrueco R, Yu J, Saucedo-Cuevas LP, Olivan M, Llobet-Navas D, Putcha P, et al. Inhibition of the autocrine IL-6-JAK2-STAT3-calprotectin axis as targeted therapy for HR-/HER2+ breast cancers. Genes Dev. 2015;29(15):1631–48.

44. Putcha P, Yu J, Rodriguez-Barrueco R, Saucedo-Cuevas L, Villagrasa P, Murga-Penas E, et al. HDAC6 activity is a non-oncogene addiction hub for inflammatory breast cancers. Breast Cancer Res. 2015;17(1):149.

45. Davoli A, Hocevar BA, Brown TL. Progression and treatment of HER2-positive breast cancer. Cancer Chemother Pharmacol. 65(4):611–23.

46. Guo L, Kozlosky CJ, Ericsson LH, Daniel TO, Cerretti DP, Johnson RS. Studies of ligand-induced site-specific phosphorylation of epidermal growth factor receptor. Journal of the American Society for Mass Spectrometry. 2003;14(9):1022–31.

47. Okamoto I. Epidermal growth factor receptor in relation to tumor development: EGFR-targeted anticancer therapy. FEBS J. 277(2):309–15.

48. Wang S, Placzek WJ, Stebbins JL, Mitra S, Noberini R, Koolpe M, et al. Novel targeted system to deliver chemotherapeutic drugs to EphA2-expressing cancer cells. J Med Chem. 2012;55(5):2427–36.

49. Birtwistle MR, Hatakeyama M, Yumoto N, Ogunnaike BA, Hoek JB, Kholodenko BN. Ligand-dependent responses of the ErbB signaling network: experimental and modeling analyses. Mol Syst Biol. 2007;3:144.

50. Rikova K, Guo A, Zeng Q, Possemato A, Yu J, Haack H, et al. Global survey of phosphotyrosine signaling identifies oncogenic kinases in lung cancer. Cell. 2007;131(6):1190–203.

51. Soda M, Choi YL, Enomoto M, Takada S, Yamashita Y, Ishikawa S, et al. Identification of the transforming EML4-ALK fusion gene in non-small-cell lung cancer. Nature. 2007;448(7153):561–6.

52. Laurent C, Do C, Gascoyne RD, Lamant L, Ysebaert L, Laurent G, et al. Anaplastic lymphoma kinase-positive diffuse large B-cell lymphoma: a rare clinicopathologic entity with poor prognosis. J Clin Oncol. 2009;27(25):4211–6.

53. Janoueix-Lerosey I, Lequin D, Brugieres L, Ribeiro A, de Pontual L, Combaret V, et al. Somatic and germline activating mutations of the ALK kinase receptor in neuroblastoma. Nature. 2008;455(7215):967–70.

54. Sokai A, Enaka M, Sokai R, Mori S, Mori S, Gunji M, et al. Pulmonary Inflammatory Myofibroblastic Tumor Harboring EML4-ALK Fusion Gene. Jpn J Clin Oncol. 2014;44(1):93–6.

55. Bender C, Henjes F, Frohlich H, Wiemann S, Korf U, Beissbarth T. Dynamic deterministic effects propagation networks: learning signalling pathways from longitudinal protein array data. Bioinformatics. 2010;26(18):i596-i602.

56. Koivunen JP, Mermel C, Zejnullahu K, Murphy C, Lifshits E, Holmes AJ, et al. EML4-ALK fusion gene and efficacy of an ALK kinase inhibitor in lung cancer. Clin Cancer Res. 2008;14(13):4275–83.

57. TCGA-Consortium. Comprehensive genomic characterization defines human glioblastoma genes and core pathways. Nature. 2008;455(7216):1061–8.

58. Crystal AS, Shaw AT. Variants on a theme: a biomarker of crizotinib response in ALK-positive non-small cell lung cancer? Clin Cancer Res. 2012;18(17):4479–81.

59. Shaw AT, Solomon B. Targeting anaplastic lymphoma kinase in lung cancer. Clin Cancer Res. 2011;17(8):2081–6.

60. . !!! INVALID CITATION !!! [24].

61. Voena C, Di Giacomo F, Panizza E, D'Amico L, Boccalatte FE, Pellegrino E, et al. The EGFR family members sustain the neoplastic phenotype of ALK+ lung adenocarcinoma via EGR1. Oncogenesis. 2013;2:e43.

62. Chan BA, Hughes BG. Targeted therapy for non-small cell lung cancer: current standards and the promise of the future. Transl Lung Cancer Res. 2015;4(1):36–54.

63. Landi L, Minuti G, D'Incecco A, Cappuzzo F. Targeting c-MET in the battle against advanced nonsmall-cell lung cancer. Curr Opin Oncol. 2013;25(2):130–6.

64. Borisy AA, Elliott PJ, Hurst NW, Lee MS, Lehar J, Price ER, et al. Systematic discovery of multicomponent therapeutics. Proc Natl Acad Sci U S A. 2003;100(13):7977–82.

65. Chou TC. Drug Combination Studies and Their Synergy Quantification Using the Chou-Talalay Method. Cancer research. 2010;70(2):440–6.

66. Zhang G, Fang B, Liu RZ, Lin H, Kinose F, Bai Y, et al. Mass spectrometry mapping of epidermal growth factor receptor phosphorylation related to oncogenic mutations and tyrosine kinase inhibitor sensitivity. J Proteome Res. 2011;10(1):305–19.

67. Valencia K, Ormazabal C, Zandueta C, Luis-Ravelo D, Anton I, Pajares MJ, et al. Inhibition of collagen receptor discoidin domain receptor-1 (DDR1) reduces cell survival, homing, and colonization in lung cancer bone metastasis. Clin Cancer Res. 2012;18(4):969–80.

68. Scott J, Ideker T, Karp RM, Sharan R. Efficient algorithms for detecting signaling pathways in protein interaction networks. Journal of computational biology : a journal of computational molecular cell biology. 2006;13(2):133–44.

69. Steffen M, Petti A, Aach J, D'Haeseleer P, Church G. Automated modelling of signal transduction networks. BMC bioinformatics. 2002;3:34.

70. Kelley BP, Sharan R, Karp RM, Sittler T, Root DE, Stockwell BR, et al. Conserved pathways within bacteria and yeast as revealed by global protein network alignment. Proceedings of the National Academy of Sciences of the United States of America. 2003;100(20):11394–9.

71. Shimoni Y, Fink MY, Choi SG, Sealfon SC. Plato's cave algorithm: inferring functional signaling networks from early gene expression shadows. PLoS Comput Biol. 2010;6(6):e1000828.

72. Budak G, Eren Ozsoy O, Aydin Son Y, Can T, Tuncbag N. Reconstruction of the temporal signaling network in Salmonella-infected human cells. Front Microbiol. 2015;6:730.

73. Zhang QC, Petrey D, Deng L, Qiang L, Shi Y, Thu CA, et al. Structure-based prediction of protein-protein interactions on a genome-wide scale. Nature. 2012;490(7421):556–60.

74. Lamb J, Crawford ED, Peck D, Modell JW, Blat IC, Wrobel MJ, et al. The Connectivity Map: using gene-expression signatures to connect small molecules, genes, and disease. Science. 2006;313(5795):1929–35.

75. Mok TS, Wu YL, Thongprasert S, Yang CH, Chu DT, Saijo N, et al. Gefitinib or carboplatin-paclitaxel in pulmonary adenocarcinoma. The New England journal of medicine. 2009;361(10):947–57.

76. Steuer R, Kurths J, Daub CO, Weise J, Selbig J. The mutual information: detecting and evaluating dependencies between variables. Bioinformatics. 2002;18 Suppl 2:S231–40.

77. Gandhi J, Zhang JL, Xie Y, Soh J, Shigematsu H, Zhang W, et al. Alterations in Genes of the EGFR Signaling Pathway and Their Relationship to EGFR Tyrosine Kinase Inhibitor Sensitivity in Lung Cancer Cell Lines. PloS one. 2009;4(2).

